# Prediction of inter-residue contacts with DeepMetaPSICOV in CASP13

**DOI:** 10.1101/586800

**Authors:** Shaun M. Kandathil, Joe G. Greener, David T. Jones

## Abstract

In this article, we describe our efforts in contact prediction in the CASP13 experiment. We employed a new deep learning-based contact prediction tool, DeepMetaPSICOV (or DMP for short), together with new methods and data sources for alignment generation. DMP evolved from MetaPSICOV and DeepCov and combines the input feature sets used by these methods as input to a deep, fully convolutional residual neural network. We also improved our method for multiple sequence alignment generation and included metagenomic sequences in the search. We discuss successes and failures of our approach and identify areas where further improvements may be possible. DMP is freely available at: https://github.com/psipred/DeepMetaPSICOV.

## 1. Introduction

The value of accurate inter-residue contact predictions in protein tertiary structure prediction is now well established. Recent years have seen marked improvements in accurate prediction of contacts, driven by improvements in methodology, most recently using meta-predictors and deep learning (Adhikari, et al., 2017; Buchan and Jones, 2018; Jones and Kandathil, 2018; Jones, et al., 2015; Liu, et al., 2018; Wang, et al., 2017; Wang, et al., 2018). For our contact prediction effort in CASP13, we developed DeepMetaPSICOV (abbreviated DMP), a contact predictor based on a deep, fully convolutional residual network and a large input feature set. DMP is a logical extension and combination of our previous methods MetaPSICOV (Buchan and Jones, 2018; Jones, et al., 2015) and DeepCov (Jones and Kandathil, 2018). The method is capable of precise predictions for a variety of proteins, including membrane proteins and those with relatively shallow sequence alignments. We also employed expanded sequence data banks for multiple sequence alignment (MSA) generation during the prediction season, which led to an overall enhancement in contact precision. In this paper, we will describe the method, its performance in CASP13, and successes and failures of our approach.

## 2. Methods

### 2.1 Feature sets

The input features to DMP comprise the sequence profile, predicted secondary structure, solvent accessibility, and other features used in MetaPSICOV (see Supplementary Information for complete details). Features defined on single residues are converted into 2D maps by striping them horizontally and vertically. Other features such as the outputs from PSICOV (Jones, et al., 2012), CCMpred (Seemayer, et al., 2014) and FreeContact (Kaján, et al., 2014) are used without modification, since they are defined on residue pairs. The 58-channel MetaPSICOV inputs are combined with the 441-channel DeepCov covariance matrices, which contain raw covariance values calculated for each pair of positions in the sequence alignment, for each pair of residue types (Jones and Kandathil, 2018). Two additional channels encode sequence separation between residue pairs and the sequence bounds; the latter is simply a channel where all input values are set to 1.

### 2.2 Model architecture

The DMP model is a deep, fully convolutional residual neural network (ResNet; Figure 1). This type of model is known to be highly performant in image recognition tasks (He, et al., 2016), as well as in contact prediction (Wang, et al., 2017; Wang, et al., 2018). In our model, the 501-channel inputs are fed to a convolutional Maxout layer (Goodfellow, et al., 2013), which reduces the input dimensionality from 501 to 64. Instance normalisation (Ulyanov, et al., 2016) is applied to the output of this layer, and the output is fed to a series of residual blocks. Each residual block (right-hand panel of Figure 1) is a set of two dilated 2D convolutional layers, each with 5×5 filters, 64 output feature maps and Rectified Linear Unit (ReLU) activation functions, together with a residual or skip-connection that adds the input of the block to its output, before passing the result through a final ReLU nonlinearity. A total of 18 residual blocks are used. Each residual block alternates between using regular and dilated 5×5 filters, with the dilation rates increasing in later residual blocks. Dilations are applied as a means to rapidly grow the receptive field of the network to encompass the whole protein input. The dilation rates used are 1, 2, 4, 8, 16, 32 and 64. After the last residual block employing dilated convolutions, a few additional blocks comprising regular (non-dilated) convolutions are used; the dilation rates used for each residual block are given in Supplementary Table S2.

**Figure 1:**
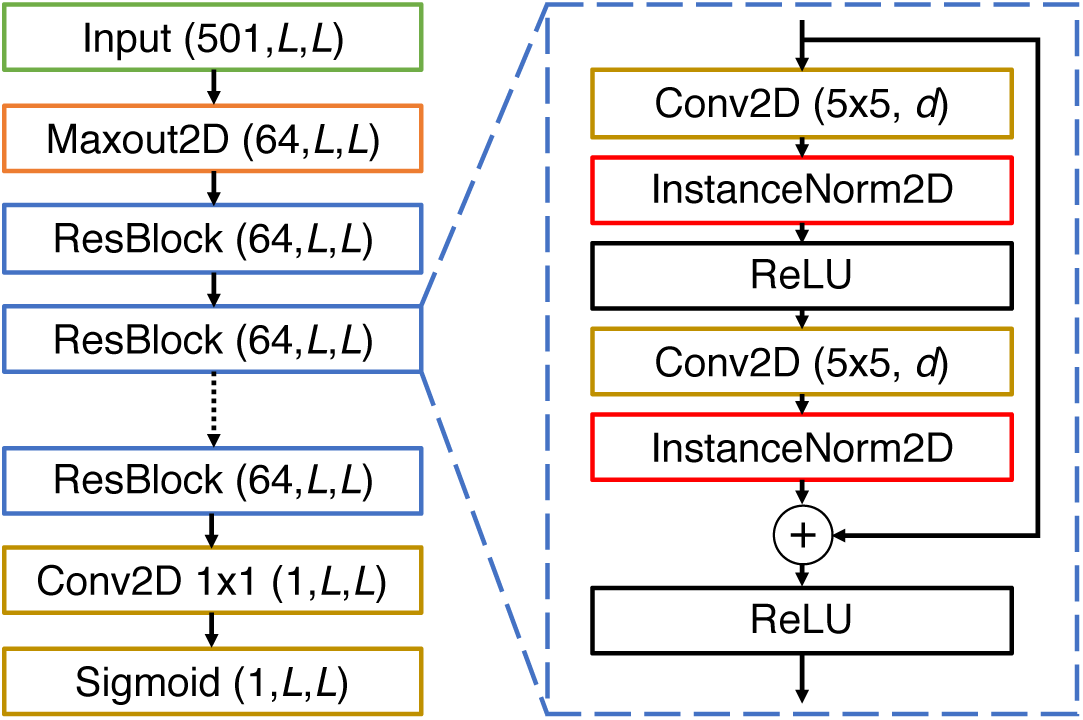
Architecture of the DeepMetaPSICOV residual neural network model. On the left, the overall organisation of the model is shown, beginning with the inputs, and ending in the final sigmoid output layer. The numbers in parentheses represent the dimensionality of the output from each layer in the format (*number of feature channels, width, height*). The network takes in input features for a protein of length *L* and produces correspondingly sized output. Most of the model is comprised of 18 residual blocks (denoted ResBlock; only a few are shown), and the structure of each block is shown on the right. The convolutional layers (Conv2D) in a residual block have 5×5 filters with a dilation rate *d*. The values of *d* for each residual block in the model are given in Supplementary Table S2.

Following the residual blocks, the output layer of the model comprises a 2D convolutional layer with a single 1×1 filter and a sigmoid nonlinearity, with instance normalisation applied before the nonlinearity. To get the predicted scores for each residue pair, we average the values predicted for residue pairs (i,j) and (j,i) as in DeepCov. The final predicted score is the average of predictions from 5 versions of the DMP model, trained on the same input data independently using different random number seeds.

### 2.3 Data augmentations

Data augmentation procedures are commonly used to improve the generalisation and robustness of models that operate on images or audio. The idea is to generate artificial, but plausible, new training examples by applying transformations to a set of “true” examples. For example, if one is interested in recognising a piece of music, one could generate new versions of a given recording by generating versions played at slightly different tempos. For contact prediction, we used three procedures inspired by techniques used in image analysis:

#### 2.3.1 Loop sampling

Loop regions in many proteins are capable of tolerating insertions and deletions without significantly affecting the overall contact pattern. Therefore, synthetic training examples can be generated by simply masking or deleting rows and columns in the input tensors corresponding to residues in loops, and by masking or deleting the corresponding sections in the contact maps as well (Figure 2a).

**Figure 2:**
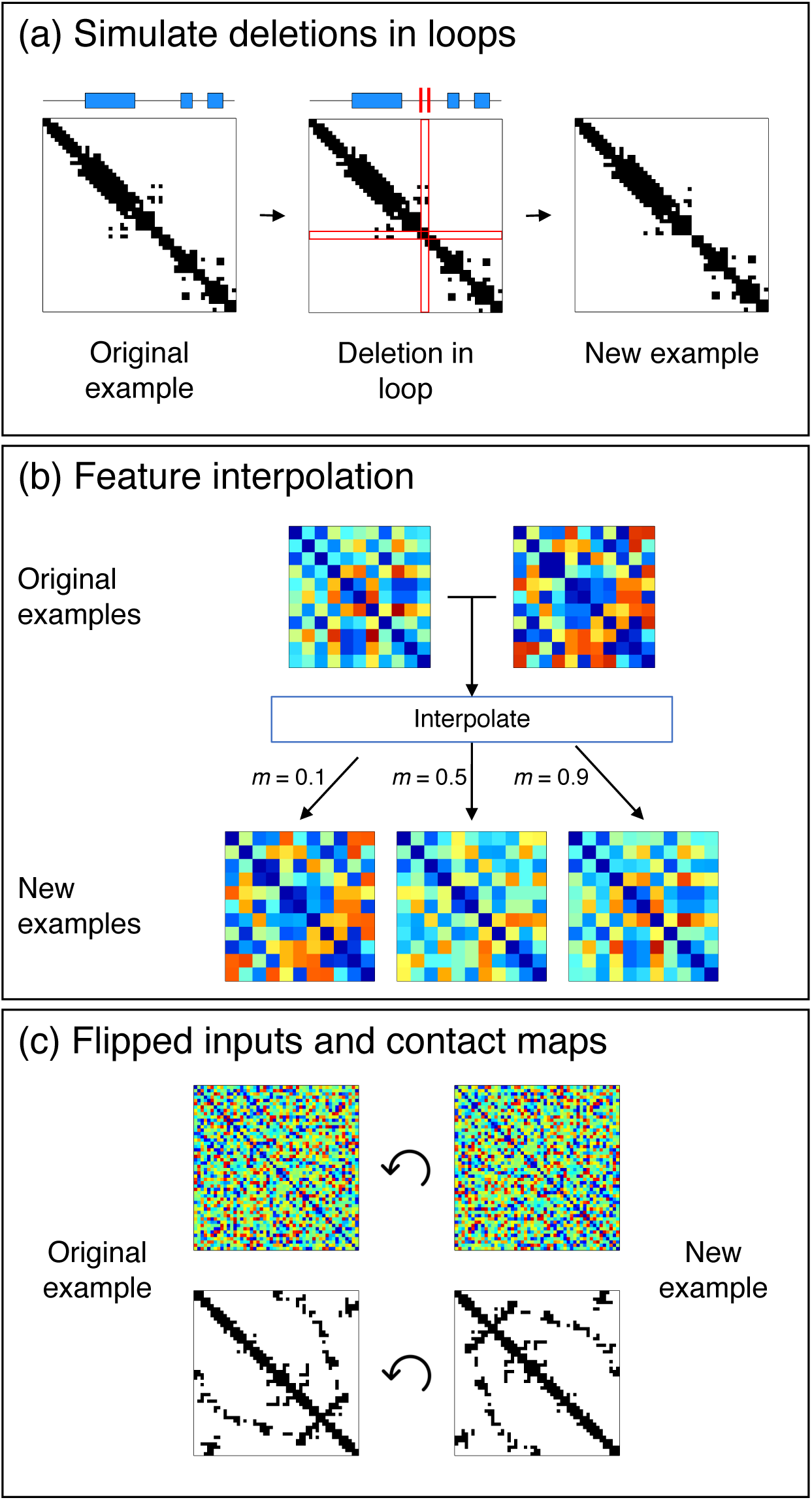
The data augmentation procedures used during the training of DeepMetaPSICOV. (a) Deletions in loops can be simulated by probabilistically removing rows and columns in the input tensors and contact maps corresponding to residues classified as loops by DSSP. The DSSP assignment for an example protein is shown above its contact map, with blue rectangles representing alpha helices, and line segments representing loops. (b) Input tensors generated using different alignments can be linearly interpolated to produce new training examples, simulating inputs generated from alignments of varying quality. Inputs thus generated for a given protein are mapped to the same contact maps. (c) New examples are generated by flipping the input feature tensors and contact maps by 180°, corresponding to a reversal of the chain direction.

Loop residues are determined according to the DSSP (Kabsch and Sander, 1983; Sander, et al., 2010) assignment for each protein in the training set. Features for residues given either no assignment or an assignment of ‘S’ corresponding to bends are considered for removal with a probability of 0.3. The corresponding rows and columns in the true contact map for the training example are also removed, and the channel encoding sequence separation (Supplementary Table S1) is also recomposed to reflect the modified sequence length. The overall procedure is applied with a probability of 0.5 and only on proteins which have 40% or fewer of their residues classified as loop according to the above definition.

#### 2.3.2 Feature interpolation

Accurate prediction of contacts is challenging when one is faced with low-quality or shallow alignments, because one obtains sparse and/or inaccurate estimates of substitution statistics. To make our method robust to alignments of lower quality, we train our models on two versions of sequence alignments: those obtained using HHblits and the pre-clustered uniprot20_2016_02 database (pre-CASP12), and those obtained using PSI-BLAST searches on the Swiss-Prot sequence database. In general, the Swiss-Prot alignments tend to be of significantly lower quality as compared to the uniprot20 alignments. As illustrated in Figure 2b, the augmentation procedure constructs synthetic training examples by linearly interpolating between two input tensors:

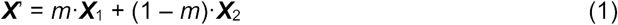

where ***X***’ is the synthetic training example, ***X***_1_ and ***X***_2_ are the original training examples, and *m* is a scalar chosen uniformly at random in the range [0, 1]. In our case, ***X***_1_ and ***X***_2_ correspond to the input feature tensors generated using uniprot20 and Swiss-Prot alignments, respectively, for a given protein in the training set. Using this procedure, we can simulate input feature tensors obtained from alignments of continuously varying quality, thus improving the model’s robustness to low-quality alignments.

The above procedure is similar to those used in the Synthetic Minority Over-sampling Technique (Chawla, et al., 2002) and the *mixup* method (Zhang, et al., 2017). The key differences relative to SMOTE and *mixup* are that (a) the interpolation in our method is not designed to over-sample any particular type of training example, and (b) interpolation is performed only on the input features; once a synthetic training example is created, it is mapped to the same (true) contact map as the original training examples.

#### 2.3.3 Flipped input feature tensors and contact maps

In image recognition, a rotated image and the original obviously contain the same information. Although contact maps cannot be arbitrarily rotated, a rotation of 180° is permitted, as this corresponds to a reversal of the protein chain direction (N and C termini are exchanged; Figure 2c). Although the resulting sequence and contact map may well not correspond to a stable, folded, and functional protein, it nonetheless describes a valid chain conformation. By reversing both the input tensors and the target contact maps in this way, the additional input/target pairs help regularise the network during training. This procedure is applied with a probability of 0.5. When applied, the flipped inputs and outputs are appended to their regular versions in a batch.

### 2.4 Training

Network weights were trained using batches of 8 training examples. The data augmentation procedures were applied on-the-fly as each batch was prepared. The implementation in PyTorch allows the training loop to accumulate weight gradients based on forward passes of individual examples. Following this, the network parameters can be updated using the gradients accumulated over each batch. With such a setup, training examples are passed through the network one at a time, removing the need for zero padding to have training examples of differing sizes in a batch.

The weights in the network were initialised using Xavier initialisation (Glorot and Bengio, 2010) with weights drawn from the uniform distribution. Network weights were optimised using the Adam method (Kingma and Ba, 2014) with an initial learning rate of 0.001. The binary cross-entropy between the predicted and true contacts was used as the loss function during training, with the loss calculated on residue pairs with sequence separation greater than 4. Training progress was monitored using the Matthews correlation coefficient (MCC) of the predictions on a separate validation set of proteins (see below). Once again, residue pairs fewer than 5 residues apart in sequence were excluded from the MCC calculation. Training was stopped when the MCC on the validation set did not improve for a number of consecutive epochs.

### 2.5 Datasets for training and testing

DMP was trained using the same set of 6729 proteins and alignments used to train DeepCov (Jones and Kandathil, 2018). The proteins in the training set were selected such that any two chains are < 25% sequence-identical and any single chain has fewer than 500 residues. Chains with missing residues were also excluded. The training set includes both single- and multi-domain proteins and has no overlap with the CASP12 free-modelling domains, which was used as a test set during development. Overlap between the training and test sets was assessed using ECOD database classification, rather than sequence identity, as the former is a much more rigorous procedure for exclusion of topologically similar proteins. Proteins were removed from the training set if they were in the same ECOD T-group as a test example. The validation set comprised the first 200 chains in the alphabetically ordered list of PDB and chain identifiers for the training set. During development, the effectiveness of the model was assessed on a variety of datasets including the CASP11 and CASP12 free-modelling (FM) domains, the PSICOV150 set (Jones, et al., 2012), and membrane proteins from Nugent and Jones (2012) and Hayat, et al. (2015).

### 2.6 Multiple sequence alignment (MSA) generation

Having a deep, diverse multiple sequence alignment for a protein of interest is essential for successful contact prediction. It has been established that metagenomic sequence collections are a rich source of sequence data that can be used for this purpose (Ovchinnikov, et al., 2017). Therefore, in CASP13 we improved upon our previous approach for generating deeper MSAs (Buchan and Jones, 2018; Kosciolek and Jones, 2016) by including both UniRef100 and metagenomic sequences in the search. Additionally, we used profile HMMs rather than single sequences to build the target-specific HHblits database. The procedure is described in detail below.

Each target sequence was used as a query for an initial HHblits (Remmert, et al., 2011) search against the UniClust30 database provided by the Söding group. If at least 10*L* raw sequences were found (where *L* is the length of the target sequence), the alignment was used as-is. For targets for which fewer than 10*L* sequences were obtained, the query sequence was scanned against a custom sequence database using jackHMMER (Clements, et al., 2011; Johnson, et al., 2010). This custom database is the set union of UniRef100 and the EBI MGnify (Tarkowska, et al., 2017) protein sequences at a sequence identity threshold of 100%. Significant hits obtained from this search were then clustered using kClust (Hauser, et al., 2013) and the clusters were aligned using MAFFT (Katoh and Standley, 2013). These alignments and the alignment from the initial HHblits search were then used to build a HHblits database specific to the target sequence. A final HHblits search was run against this target-specific HHblits database to derive the final MSA.

### 2.7 Calculation of effective sequence count (*M*_*eff*_)

Sequences in the MSA for each target were clustered using CD-HIT (Fu, et al., 2012; Li and Godzik, 2006) at a sequence identity threshold of 62% and a word size of 4. The number of clusters returned by CD-HIT was taken as the *M*_*eff*_. Unless otherwise mentioned, *M*_*eff*_ values are calculated on the alignment obtained by the MSA generation procedure described above for the full-length target sequence.

### 2.8 Automatic domain parsing

We attempted to automatically parse domains in each target sequence using the same approach we used in CASP12. Briefly, each target sequence was first run through the alignment generation and contact prediction steps to generate an initial contact list. Using HHblits, the target sequence was scanned against the PDB70 database provided by the Söding group. Regions of the sequence that did not match a PDB template and that were at least 30 residues long were extracted, and the alignment generation and contact prediction steps were re-run on the putative domain sequence. The contact scores predicted for such domains were then copied back into the relevant region(s) of the initial contact list to yield the final prediction.

## 3. Results

### 3.1 Performance in CASP13

Our move to a deep residual neural network model for generating contact predictions proved to be quite successful. In an early test on the CASP12 FM domains, we observed that DMP was substantially more precise than MetaPSICOV2 and DeepCov on the same input alignments. Addition of the data augmentation procedures and averaging predictions over 5 versions of the trained model also led to small improvements in mean precision on these targets.

Table 1 shows the precision obtained by DMP on the domains classified as FM or FM/TBM by the CASP13 assessors. Over these targets, DMP obtained a mean precision of 66.18% when considering the top-L/5 long-range contacts. Our predictions were more than 90% precise for 16 domains, and a top-L/5 precision of 100% was achieved on 7 of these domains. Notably, some very precise predictions were obtained even though the MSA for the target had a low effective sequence count; considering the 16 domains in Table 1 for which our alignments had an *M*_*eff*_ <= 50, DMP obtained a mean long-range precision of 44.48% for the top-L/2 contacts, and 57.88% on the top-L/5 contacts. The corresponding mean precision values considering both medium and long-range contacts are 61.11% and 76.33%. This represents a strong improvement in our ability to accurately predict contacts for relatively shallow MSAs, especially when one considers (for example) that PSICOV requires many hundreds of effective sequences in the MSA to achieve similar precision (Jones, et al., 2012).

**Table 1:** Performance of DMP in CASP13. Top-L/5 long-range precision is shown for 43 FM and FM/TBM domains. Targets are ordered by domain classification, followed by domain identifier. *M*_*eff*_ values (see Section 2.7) are calculated on the MSA for the full-length target sequence, and so different domains of the same target have the same *M*_*eff*_.

| Domain | Classification | Length | Precision (%) | $M_{eff}$ |
| --- | --- | --- | --- | --- |
| T0950-D1 | FM | 342 | 94.20 | 111 |
| T0953s2-D2 | FM | 111 | 100.00 | 180 |
| T0953s2-D3 | FM | 93 | 93.75 | 180 |
| T0957s1-D1 | FM | 108 | 36.36 | 43 |
| T0957s2-D1 | FM | 155 | 87.10 | 37 |
| T0960-D2 | FM | 84 | 17.65 | 70 |
| T0963-D2 | FM | 82 | 11.76 | 58 |
| T0968s1-D1 | FM | 119 | 54.17 | 116 |
| T0968s2-D1 | FM | 116 | 39.13 | 229 |
| T0969-D1 | FM | 354 | 98.59 | 645 |
| T0975-D1 | FM | 293 | 80.70 | 4918 |
| T0980s1-D1 | FM | 105 | 100.00 | 50 |
| T0981-D2 | FM | 80 | 31.25 | 6 |
| T0986s2-D1 | FM | 155 | 80.65 | 56 |
| T0987-D1 | FM | 185 | 100.00 | 23 |
| T0987-D2 | FM | 207 | 92.50 | 23 |
| T0989-D1 | FM | 134 | 55.56 | 65 |
| T0989-D2 | FM | 112 | 43.48 | 65 |
| T0990-D1 | FM | 76 | 37.50 | 31 |
| T0990-D2 | FM | 231 | 36.17 | 31 |
| T0990-D3 | FM | 213 | 55.81 | 31 |
| T0991-D1 | FM | 111 | 0.00 | 1 |
| T0998-D1 | FM | 166 | 44.12 | 8 |
| T1000-D2 | FM | 431 | 95.95 | 873 |
| T1001-D1 | FM | 139 | 10.71 | 11 |
| T1010-D1 | FM | 210 | 88.10 | 89 |
| T1015s1-D1 | FM | 88 | 27.78 | 580 |
| T1017s2-D1 | FM | 128 | 72.00 | 87 |
| T1021s3-D1 | FM | 178 | 94.12 | 979 |
| T1021s3-D2 | FM | 101 | 15.00 | 979 |
| T1022s1-D1 | FM | 156 | 90.62 | 1393 |
| T0949-D1 | FM/TBM | 139 | 100.00 | 6067 |
| T0953s2-D1 | FM/TBM | 44 | 66.67 | 180 |
| T0958-D1 | FM/TBM | 77 | 100.00 | 22 |
| T0970-D1 | FM/TBM | 97 | 88.24 | 43 |
| T0978-D1 | FM/TBM | 413 | 87.80 | 1602 |
| T0981-D3 | FM/TBM | 203 | 100.00 | 6 |
| **T0986s1-D1** | FM/TBM | 92 | 73.68 | 187 |
| **T0992-D1** | FM/TBM | 107 | 100.00 | 436 |
| **T0997-D1** | FM/TBM | 185 | 94.59 | 963 |
| **T1005-D1** | FM/TBM | 326 | 93.94 | 872 |
| **T1008-D1** | FM/TBM | 77 | 6.25 | 5 |
| **T1019s1-D1** | FM/TBM | 58 | 50.00 | 267 |

### 3.2 Successes and failures in MSA generation

The addition of metagenomic sequences to our MSA generation step proved beneficial in an initial test on a subset of the CASP12 FM domains, where we found that we were able to obtain as many as double the number of sequences as compared to using UniRef100 alone. In CASP13, we saw an improvement in alignment depth over using HHblits alone (Figure 3a) for all but 3 targets which had fewer than 10L raw sequences in the initial HHblits MSA. The new procedure for MSA generation guarantees that the MSA derived by searching the custom database of UniRef100 + EBI MGnify sequences will have an equal or greater number of (raw) sequences as compared to the initial HHblits MSA, and thus all points in Figure 3a are on or above the dashed line. The increase in alignment depth translated into more precise contact predictions overall (Figure 3b). Strong improvements in precision were seen when using the deeper MSAs on domains T0958-D1, T1010-D1, and T0957s2-D1, among several others.

**Figure 3:**
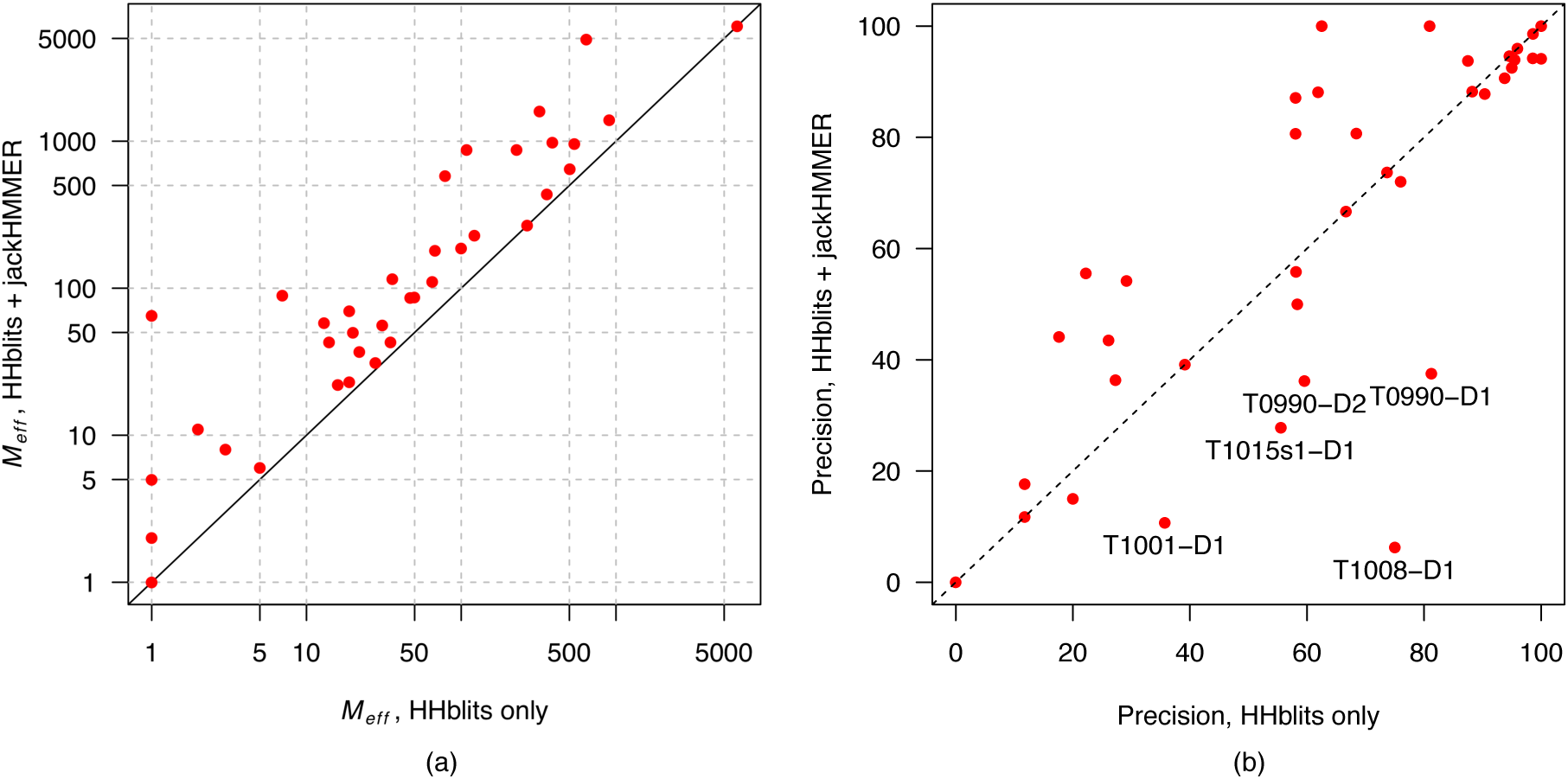
(a) Comparison of effective sequence count (*M*_*eff*_) between alignments generated using only HHblits, or HHblits and jackHMMER. In the latter case, the jackHMMER search makes use of UniRef100 and EBI MGnify metagenomic protein sequences. (b) Plot of top-L/5 long-range precision values obtained using the deeper alignments versus those obtained using HHblits only. Using the deeper alignments was beneficial overall, although there are a few domains for which just the HHblits alignment would have provided much higher precision; these are marked.

From Figure 3b, there are a few cases in which deeper alignments led to significantly reduced contact precision. Reduced performance with deeper alignments could indicate (among other factors) misalignment, ‘blurring’ or loss of structural signal in MSAs with very distant sequence relatives, or that the MSA contains sequences incorrectly matched due to profile drift. We found evidence of the latter on target T1015s1-D1, for which we obtained a top-L/5 long-range precision of 27.78%. The full MSA (*M*_*eff*_ = 580) for this target shows very highly conserved CXC and CXXC motifs, corresponding to a metal binding site in the tertiary structure. Despite these patterns of conservation, many of the sequences in the alignment appear to be artefactual hits brought in by profile drift. Indeed, when predicting contacts using just the initial HHblits MSA (*M*_*eff*_ = 79), the top-L/5 long-range precision jumps to 55.56%. These observations highlight challenges encountered when using a one-size-fits-all approach to MSA generation, and this is an area that we plan to develop further.

### 3.3 Domain parsing

Our automatic domain parsing procedure detected domains on a total of 20 out of the 90 regular targets during the prediction season. Of the contact prediction targets, our domain parsing procedure detected domains for 5 targets corresponding to 6 domains (Table 2). Of these, only T0981-D3 benefitted clearly from the automated domain parsing, gaining between 35 (top-L) and 12.2 (top-L/5) percentage points in precision. T0981-D2 showed a mixed result, gaining significantly in terms of top-L/10 precision, but showing no difference or worse precision on longer contact lists. No change in precision is obtained on T0949, reflecting the fact that the alignment for the full-length sequence was already very deep (Table 1). In summary, from a contact prediction perspective, automatic domain parsing results in little or no benefit in terms of contact precision when domains are detected. These findings are in general agreement with our findings in CASP12 (Buchan and Jones, 2018), where we observed only minor improvements in top-L/5 contact precision on a few targets after parsing domains.

**Table 2:** Change in precision after automatic domain parsing. Values are expressed as percentage point differences relative to predictions made without domain parsing.

| **ΔPrecision after domain parsing (percentage points)** |  |  |  |  |
| --- | --- | --- | --- | --- |
| **Domain** | **Top-L** | **Top-L/2** | **Top-L/5** | **Top-L/10** |
| **T0949-D1** | 0.00 | 0.00 | 0.00 | 0.00 |
| **T0960-D2** | 0.00 | 4.76 | 0.00 | 0.00 |
| **T0978-D1** | 4.60 | 3.38 | 3.61 | -4.76 |
| **T0981-D2** | -6.25 | -12.50 | 0.00 | 37.50 |
| **T0981-D3** | 34.98 | 26.47 | 12.20 | 0.00 |
| **T1000-D2** | 0.54 | 1.09 | 0.00 | 0.00 |

Non-detection of domains proved to be a significant issue for some targets. A clear example of this was T1021s3, for which our pipeline did not detect any domains. The MSA for this target had 3112 raw sequences (*M*_*eff*_ = 979). Figure 4 shows the gap fraction in each column of the MSA generated for this target, along with the official domain boundaries. The region of this MSA corresponding to the second domain is almost entirely covered by gaps, meaning there is little information to use. Consequently, the contact precision in this domain is very low. In contrast, contacts in the first domain are very precise owing to much better coverage in this region and the high effective sequence count of the alignment. In this instance, the gap fraction in the MSA columns would have alerted us to the existence of the second domain, however gap content on its own is very unlikely to be a general solution to the problem.

**Figure 4:**
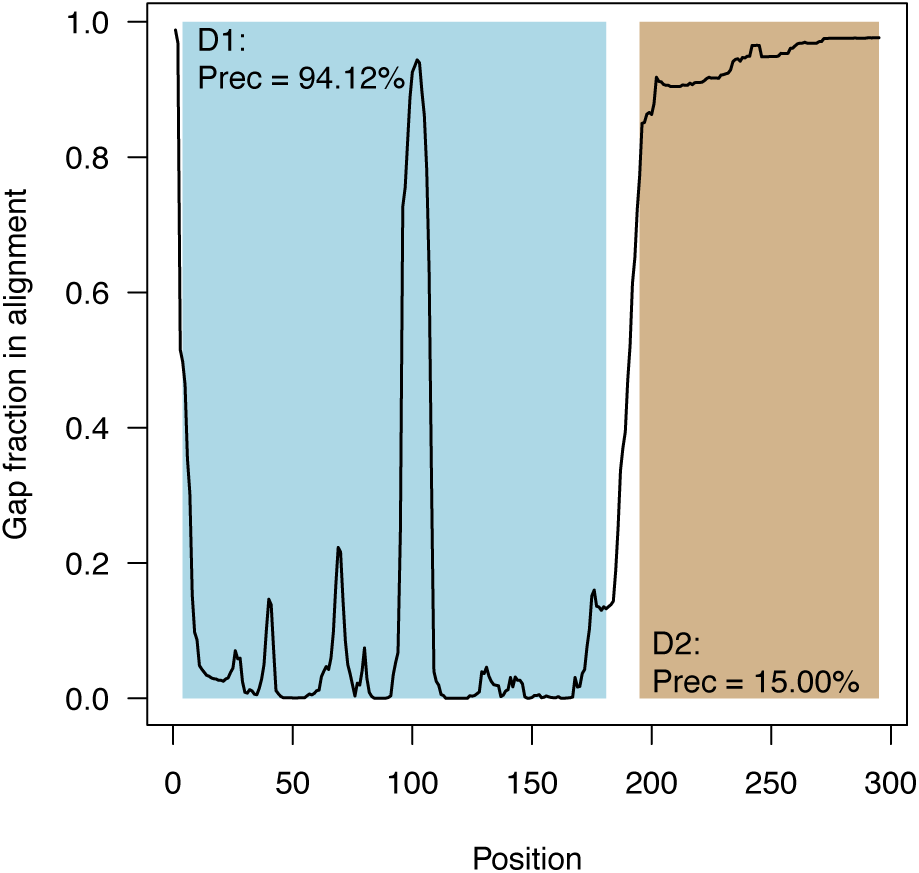
Gap fraction per column in the MSA generated for target T1021s3 (3112 raw sequences, *M*_*eff*_ = 979). Official domain boundaries are shaded in light blue and brown, and the precision obtained by DMP on these domains (long-range, top-L/5) is shown. The region of the MSA covering the C-terminal domain D2 is comprised mostly of gaps, and thus has little to no information content. Consequently, the obtained contact precision on this domain is much lower than that obtained for D1.

### 3.4 Incorrect calculation of mutual information

After the prediction season, we noticed that our calculation of mutual information (MI) values during inference was incorrect due to a bug in an in-house program. This bug affected all our predictions during the CASP13 prediction season, although we verified that it did not affect training of the DMP models. After correcting the bug, we determined its impact on performance by repeating our predictions on the 43 domains in Table 1. As with our “official” predictions, contacts were predicted for full-chain sequences, and precision was assessed on the official domains for these targets.

Incorrect MI calculations led to a loss of roughly 2-4% mean long-range precision on this set of domains, depending on the length of the contact list considered (Table 3). The worst-affected cases were domains T0998-D1, T0990-D2 and T1001-D1, for which using the correct version results in gains of 20.59, 21.28 and 39.29 percentage points in top-L/5 precision respectively relative to the incorrect version. Interestingly, incorrect MI values tend to have a greater impact on contact precision for targets with an *M*_*eff*_ of around 100 or lower (Figure 5). This observation suggests that MI features may have a greater influence on the predictions made by the DMP neural network model when other features are sparse and appear to be an important contributor to performance on MSAs with low *M*_*eff*_.

**Table 3:** Mean precision values obtained for 43 CASP13 domains using correct or incorrect MI values in the input to the DMP neural network model. The ‘Incorrect MI’ results were obtained during the CASP13 prediction season due to a bug in our MI calculations, whereas the ‘Correct MI’ data were obtained post-hoc using corrected MI values and operating on the same inputs.

| **Mean long-range precision (%)** |  |  |  |  |
| --- | --- | --- | --- | --- |
|  | **Top-L** | **Top-L/2** | **Top-L/5** | **Top-L/10** |
| **Correct MI** | 43.87 | 56.94 | 69.54 | 75.84 |
| **Incorrect MI** | 41.93 | 53.49 | 66.27 | 71.25 |

**Figure 5:**
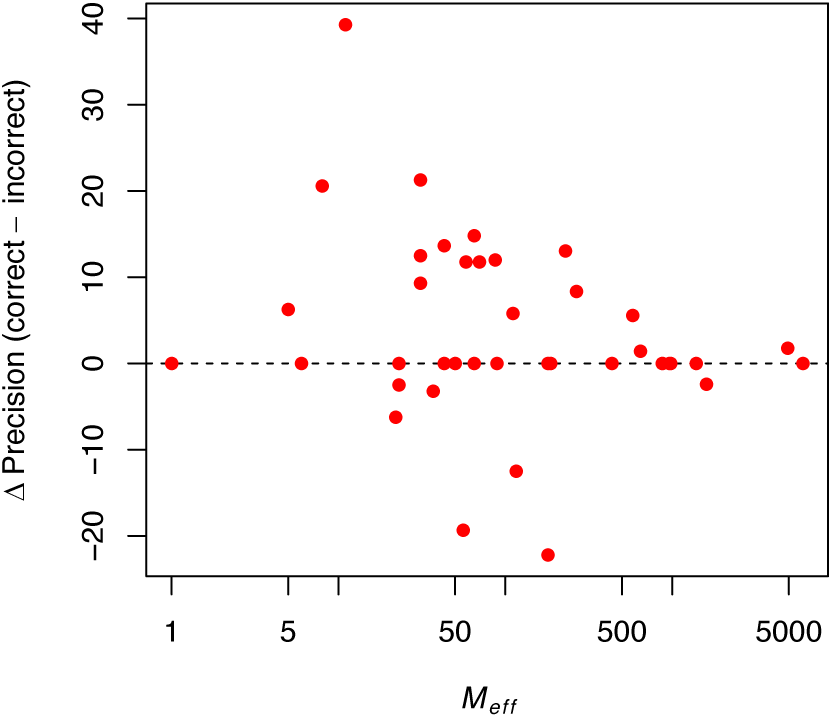
Impact of incorrect mutual information (MI) calculations on top-L/5 long-range contact precision. Values are expressed as percentage point differences, with positive values indicating a gain in precision upon using the correct MI calculation.

## 4. Discussion

It is evident from our results in CASP13 (and those of other groups) that methods based on deep learning now represent the state of the art in inter-residue contact prediction. DMP is our most effective contact prediction method to date. Nevertheless, results from this CASP indicate that there is considerable room for improvement.

The addition of metagenomic sequences during MSA generation was beneficial overall, and we plan to integrate additional sources of sequence data in the future. However, in some cases the deeper alignments did not yield benefits in contact precision, and thus care must be taken that sensitive, iterated sequence homology searching procedures do not pull in unrelated sequences due to profile drift. Nevertheless, there are early indications that careful application of remote homology searching can yield even greater benefits than we were able to realise in CASP13. Towards the end of the prediction season, we experimented with an iterated version of our MSA generation procedure (SSection 2.6) which uses *hmmbuild* and *hmmsearch* instead of jackHMMER to search the custom sequence database. The advantage of this setup is that it allows the entire process of searching the custom database to be iterated using the MSA generated at the end of each round. Initial testing indicated that the procedure was prone to profile drift, and we deemed it too unstable to use as our default MSA generation strategy. However, in at least one case (T1010-D1), this procedure provided a much deeper alignment after 3 iterations (*M*_*eff*_ increased from 89 to 200), concomitant with an increase in top-L/2 medium+long range precision from 79.05% to 91.43%. Despite these encouraging results, it is not yet clear if such procedures can be reliably used in a fully automated manner, although this is something that we are keen to explore.

Further improvements in predictive accuracy may be possible by testing different architectures for the DMP neural network. Early results indicate that moving to an even deeper network architecture is beneficial, albeit with diminishing returns as network depth increases. More broadly, from the perspective of 3D structure determination, it is becoming clear that deep learning models like ours can also be used to extract much richer forms of structural information such as interatomic distances (e.g. Wang, et al., 2018; Xu, 2018). Our tertiary structure prediction effort in CASP13 did not make use of the contacts predicted by DMP. Instead, we developed a tertiary structure prediction method that uses distances predicted from the same input features used by DMP (Greener, et al., 2018), in common with the approach taken by other groups in CASP13. Initial results were encouraging, and we are continuing to develop the method.

## Acknowledgements

We are grateful to members of the group for helpful comments and discussions. We are grateful to the Söding group for quickly correcting problems with the PDB70 database during the prediction season.

## Funding

This work was supported by the Francis Crick Institute, which receives its core funding from Cancer Research UK (FC001002), the UK Medical Research Council (FC001002) and the Wellcome Trust (FC001002). This work was also supported by the European Research Council Advanced Grant ‘ProCovar’ (Project ID 695558).

