## Supplementary Information for "Prediction of inter-residue contacts with DeepMetaPSICOV in CASP13"

Table S1. Listing of all input features and their contributions to the DMP input feature tensor. For features defined on single residues, the feature values are striped horizontally and vertically to convert them into 2D feature maps with spatial dimensions of $L \times L$, where $L$ is the length of the target sequence. This causes such features to occupy twice the number of channels in the input tensor as compared to features defined on residue pairs.

| Feature | Feature defined for single residues (1) or residue pairs (2) | Dimensionality per residue or residue pair | Channels occupied in input tensor |
|---|---|---|---|
| PSIBLAST Sequence profile | 1 | 21 | 42 |
| MI | 2 | 1 | 1 |
| MIp | 2 | 1 | 1 |
| Mean contact potential | 2 | 1 | 1 |
| PSICOV contact scores | 2 | 1 | 1 |
| FreeContact (mfDCA) contact scores | 2 | 1 | 1 |
| CCMpred (plmDCA) contact scores | 2 | 1 | 1 |
| PSIPRED secondary structure | 1 | 3 | 6 |
| Shannon entropy in MSA columns | 1 | 1 | 2 |
| SOLVPRED solvent accessibility | 1 | 1 | 2 |
| log(1 + sequence separation) | 2 | 1 | 1 |
| Sequence bounds (channel of ones) | 2 | 1 | 1 |
| DeepCov raw covariances | 2 | 441 | 441 |
| | | **Total** | 501 |

Table S2. List of dilation rates *d* for each of the 18 residual blocks in the DMP ResNet. A dilation rate of *d* = 1 produces regular, non-dilated convolutions.

| Residual block | 1 | 2 | 3 | 4 | 5 | 6 | 7 | 8 | 9 | 10 | 11 | 12 | 13 | 14 | 15 | 16 | 17 | 18 |
|---|---|---|---|---|---|---|---|---|---|---|---|---|---|---|---|---|---|---|
| Dilation rate *d* | 1 | 2 | 1 | 4 | 1 | 8 | 1 | 16 | 1 | 32 | 1 | 64 | 1 | 1 | 1 | 1 | 1 | 1 |
